# Approximating the coalescent under facultative sex

**DOI:** 10.1101/846568

**Authors:** Matthew Hartfield

## Abstract

Genome studies of facultative sexual species, which can either reproduce sexually or asexually, are providing insight into the evolutionary consequences of mixed reproductive modes. It is currently unclear to what extent the evolutionary history of facultative sexuals’ genomes can be approximated by the standard coalescent, and if a coalescent effective population size *N*_*e*_ exists. Here, I determine if and when these approximations can be made. When sex is frequent (occurring at a frequency much greater than 1*/N* per reproduction per generation, for *N* the actual population size), the underlying genealogy can be approximated by the standard coalescent, with a coalescent *N*_*e*_ ≈ *N*. When sex is very rare (at frequency much lower than 1*/N*), approximations for the pairwise coalescent time can be obtained, which is strongly influenced by the frequencies of sex and mitotic gene conversion, rather than *N*. However, these terms do not translate into a coalescent *N*_*e*_. These results are used to discuss the best sampling strategies for investigating the evolutionary history of facultative sexual species.

## Introduction

Facultative sex, where individuals can either reproduce sexually or asexually, is pervasive in nature (Hartfield 2016). By switching reproduction, it is assumed that these organisms can reap the benefits of both modes (e.g., through shuffling genotypes and increasing fecundity, respectively). Genome sequence data from these organisms are being studied to determine the evolutionary consequences of mixed reproductive modes (Hartfield 2016; Nieuwenhuis and James 2016; Ho *et al*. 2019). However, difficulties arise when analysing genomes from facultative sexuals as the majority of theoretical and computational genomic methods assume obligate sex. Analyses of genome data from facultative sexuals will be aided if it is determined when general population genetic models can be used to investigate the evolutionary history of these species.

The determinants of genetic diversity in facultative sexual organisms has been analysed using coalescent theory (Kingman 1982; Wakeley 2009). In the earliest models (Brookfield 1992; Burt *et al*. 1996; Bengtsson 2003; Ceplitis 2003), offspring were either produced via parthenogenesis or sexual reproduction. Parthenogenesis limits the extent that haplotypes segregate among individual lineages. Under rare sex (occurring with frequency at most on the order of 1*/N* per reproduction per generation, for *N* the population size, hereafter denoted *𝒪*(1*/N*)), the mean coalescent time for two alleles taken from the same locus within the same individual is longer than that for two alleles taken from different individuals. Elevated within– individual coalescent times leads to “allelic sequence divergence” that raises heterozygosity (Mark Welch and Meselson 2000; Butlin 2002). Hartfield *et al*. (2016) subsequently introduced mitotic gene conversion into the model, which increases the frequency of within–individual coalescence, reducing the within–individual coalescent time (and resulting diversity) to lower than that in equivalent obligate sexuals. Given how pervasive gene conversion is during mitotic recombination (LaFave and Sekelsky 2009; Lee *et al*. 2009), then it can be a potent force in removing diversity in facultative sexuals.

Less attention has been given to determining if and when genealogies under facultative sex can be captured by the standard coalescent (Kingman 1982). If these approximations are available, they are attractive because the myriad models developed using the standard coalescent can then be applied to facultative sexuals. These approximations usually arise after specifying an appropriate effective population size *N*_*e*_. Initially defined by Wright (1931), *N*_*e*_ is defined as the population size needed for some aspect of a model to match a corresponding Wright–Fisher model with the same size. *N*_*e*_ influences many aspects of genetic evolution, including both the rate at which neutral alleles are lost by genetic drift (Fisher 1930; Wright 1931) and new alleles are introduced by mutation (Watterson 1975). In addition, for an allele with selection coefficient *s*, the efficacy of natural selection acting on it is determined by *N*_*e*_*s* (Kimura 1971).

*The N*_*e*_ of a population has been defined in several ways. Previous definitions include those based on the maximum non–unit eigenvalue of the model’s transition matrix (the ‘eigenvalue’ *N*_*e*_); the probability that two alleles are identical by descent (the ‘inbreeding’ *N*_*e*_); or the variance in allele frequencies (the ‘variance’ *N*_*e*_) (Whitlock and Barton 1997; Ewens 2004; Charlesworth and Charlesworth 2010). A more recent definition that has gained interest is the ‘coalescent effective population size’ (Whitlock and Barton 1997; Laporte and Charlesworth 2002; Sjödin *et al*. 2005). For a neutral Wright–Fisher model of size *aN* (where *a* = 1 for haploids and *a* = 2 for hermaphrodite diploids), the genealogy of a sample of *n* alleles converges to the standard coalescent if time is rescaled by *aN*. For non– standard coalescent models, if the genealogy converges to the standard coalescent after rescaling time by *aN*, but the coalescent time is scaled by a factor *c*, then the coalescent *N*_*e*_ = *N/c* (Sjödin *et al*. 2005). The coalescent *N*_*e*_ has the advantage of being relatable to the genome data being analysed, as the underlying genealogy shapes observed genetic diversity (Sjödin *et al*. 2005). Coalescent *N*_*e*_ values have been obtained in the cases of self–fertilisation (Nordborg and Donnelly 1997; Nordborg and Krone 2002), seed banks (Kaj *et al*. 2001), autotetraploids (Arnold *et al*. 2012), fluctuating population sizes (Sjödin *et al*. 2005), unequal sex–ratios (Wakeley 2009), and various models of population structure (Wakeley 2004; Sjödin *et al*. 2005; Wakeley 2009). If a coalescent *N*_*e*_ can be defined, then existing tools for genome inference based on the coalescent can be applied to genome data from facultative sexuals. In some cases, the coalescent *N*_*e*_ depends on the size of specific parameters. For example, the coalescent *N*_*e*_ with a fluctuating population size depends on how fast fluctuations occur compared to coalescent times (Sjödin *et al*. 2005).

Previous research has elucidated the neutral forces affecting *N*_*e*_ under facultative sex. Orive (1993) determined that the prevalence of multiple asexual stages before the onset of sexual reproduction tended to reduce *N*_*e*_ (similar results were obtained by Berg and Lascoux (2000)). Conversely, Balloux *et al*. (2003) demonstrated that low occurrences of sex inflate *N*_*e*_ as measured among different alleles, but decrease *N*_*e*_ as measured over genotypes. Increased variance in asexual and sexual reproductive output can further raise some measures of *N*_*e*_ (Yonezawa *et al*. 2004). However, it is unclear how mitotic gene conversion affects *N*_*e*_, or whether these previously–defined *N*_*e*_ values constitute a coalescent *N*_*e*_.

Hartfield *et al*. (2016) used a separation–of–timescale argument to show how sex and mitotic gene conversion affect coalescent times, if they acted on the same timescale as coalescent events. As these effects would shape diversity both between and within individuals on the same timescale, then the population’s genetic history cannot be captured by a single coalescent *N*_*e*_. However, this argument only covers a special case of the coalescent process. Here, I extend these separation–of–timescale arguments to show that a coalescent *N*_*e*_ can be defined if sex is very frequent (acting with probability much greater than 1*/N*). If sex is very rare (acting with probability much less than 1*/N*), it is possible to define an average pairwise time to the common ancestor that can be related to the standard coalescent, but a coalescent *N*_*e*_ cannot be defined. I will subsequently describe how the coalescent process with very rare sex can be approximated with an arbitrary number of alleles.

## Methods

### Using Möhle’s theorem to determine coalescent *N*_*e*_

The standard coalescent assumes that alleles are exchangeable (Cannings 1974; Kingman 1982), where ‘allele’ denotes a contiguous stretch of DNA sequence with negligible recombination (Nordborg and Donnelly 1997). The exchangeability assumption implies that it does not matter whether alleles are sampled from the same or different individuals. After rescaling time by the total effective number of alleles in the population *aN*_*e*_, each pair of alleles in a sample of size *n* coalesces independently, so the total rate of coalescence is 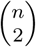 (Table 1 outlines notation used in this analysis). Hence the time between coalescent events is exponentially distributed with rate 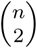, which equals 1 for *n* = 2. Under facultative sex, the exchangeability assumption breaks down and the coalescent is instead modelled using a Markov chain (Hartfield *et al*. 2016). In particular, the genealogical history of two alleles differ if they are sampled from distinct individuals (hereafter ‘unpaired’ alleles), or if two different alleles are sampled from the same locus within the same individual (hereafter ‘paired’ alleles). However, over longer timescales, it may be the case that alleles coalesce at a steady rate. In this case, a coalescent *N*_*e*_ can be inferred by rescaling time so that coalescent events occur at the same rate as in the standard coalescent (see Nordborg and Krone (2002); Sjödin *et al*. (2005) for more formal definitions).

**Table 1.** Glossary of Notation.

| Symbol | Usage |
| --- | --- |
| $a$ | Ploidy level of the population<br>($a = 1$ for haploids, $a = 2$ for hermaphrodite diploids) |
| $N$ | Actual population size (with $2N$ haplotypes for $a = 2$) |
| $N_e$ | Effective population size |
| $n$ | Number of sampled alleles in the coalescent process |
| $\sigma$ | Proportion of offspring that are produced from sexual reproduction |
| $\gamma$ | Probability of within-individual coalescence via mitotic gene conversion |
| $F$ | Inbreeding coefficient |
| $\mathbb{T}$ | Transition matrix over one generation |
| $\mathbb{A}$ | Events in $\mathbb{T}$ that are $\mathcal{O}(1)$ |
| $\mathbb{B}$ | Events in $\mathbb{T}$ that are $\mathcal{O}(\epsilon)$ for $\epsilon \ll 1$ |
| $\mathbb{P}$ | ‘Short-term’ outcomes that act over $\mathcal{O}(1)$ generations |
| $\mathbb{G}$ | ‘Long-term’ outcomes that act over $\mathcal{O}(1/\epsilon)$ generations |
| $\Omega$ | Scaled rate of sex, $2N\sigma$ |
| $\Gamma$ | Scaled rate of mitotic gene conversion, $2N\gamma$ |
| $\lambda$ | Probability of either sex or gene conversion occurring, $\lambda = \sigma + \gamma$ |
| $\phi$ | Ratio of sex to gene conversion, $\phi = \sigma/\gamma$ |
| $\Lambda$ | Scaled total probability of an event, $\Lambda = N^2\lambda$ |
| $\mathbb{E}[T_b], \mathbb{E}[T_w]$ | Expected between (within) individual coalescent time for two alleles |
| $\mathbb{E}[\tau_b], \mathbb{E}[\tau_w]$ | Expected coalescent times on the coalescent timescale<br>(scaled by $2N$ generations) |

Möhle’s theorem (Möhle 1998) is often used to separate events over short and long timescales, to determine whether a coalescent *N*_*e*_ exists. Let 𝕋 represent the discrete–time transition matrix of a structured coalescent process over one generation. Further assume that 𝕋 can be decomposed into the sum 𝕋 = 𝔸 + 𝔹*/N* + *o*(1*/N*), where *N* is the total population size. This decomposition assumes that matrices 𝔸 and 𝔹 exist as the population size becomes large (technically, as *N* → ∞); specifically, 𝔸 = 𝕋 and 𝔹 = *N* (𝕋 − 𝔸). *o*(1*/N*) are terms that approach zero faster than 1*/N* (Wakeley 2009).

Möhle (1998) proved that, if 𝕋 can be written in this manner, then the coalescent process may be described by a continuous–time rate matrix Π*(τ*) = ℙ*e*^−*τ*𝔾^, where 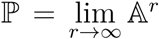 and 𝔾 = ℙ𝔹ℙ. ℙ represents ‘short–term’ events that occur on timescales much shorter than *N* generations, which describe an initial adjustment to alleles in the recent past. 𝔾 represents ‘long–term’ events that occur on 𝒪(*N*) generations. Time can then be rescaled so that coalescence events occur at the same long–term rate as in the standard model; coalescent *N*_*e*_ is subsequently inferred from this rescaling.

Möhle’s theorem can also be invoked for any scaling parameter *ϵ* ≪ 1 by writing 𝕋 = 𝔸 + *ϵ*𝔹 + *o*(*ϵ*), and as *ϵ* → 0 𝔸 = 𝕋 and 𝔹 = (𝕋 − 𝔸)*/ ϵ*. 𝔾 then represents events that occur at timescales on 𝒪(1*/ϵ*). Most application of Möhle’s theorem take *ϵ* = *c/N* for some constant *c*, which is used in the ‘frequent sex’ regime below. However, *ϵ* can also depend on other parameters, and is used in the ‘very rare sex’ case where it will be assumed that the probabilities of sex and gene conversion are both small.

### The facultative sex coalescent

The facultative sex coalescent acts in a diploid population of size *N* (i.e., there are 2*N* total alleles). Alleles are sampled from individuals, and their genealogical history is traced backwards in time. Each sampled individual can reproduce both sexually and asexually; sexual reproduction occurs with probability *σ*, and asexually (parthenogenetically) with probability 1 − *σ*. If sex occurs then each allele in an individual is inherited from two random parents sampled with replacement, so a single individual can act as both parents. Otherwise, both alleles are inherited in state from the same parent. Self–fertilisation can also be included in the model (Hartfield *et al*. 2016), but is not considered here. Mitotic gene conversion (hereafter ‘gene conversion’) acts with probability *γ*, which causes two alleles that reside within the same individual to coalesce. Note that the usage here indicates the probability of gene conversion acting per individual; it is twice the probability of gene conversion affecting a single site, as there are two possible donor strands where it can initiate (Hartfield *et al*. 2018).

Two sampled alleles can lie in one of three states: (i) they lie in different individuals, (ii) they lie in the same individual, but have not coalesced, (iii) they have coalesced. The coalescent history can be determined by the following transition matrix (Hartfield *et al*. 2016, Eq. 10 without self–fertilisation):

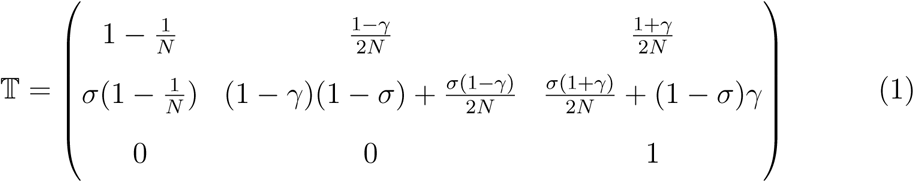

Figure 1 illustrates how the transition probabilities are determined. Row 1 describes transitions from the state where alleles reside in different individuals. Going back one generation, two alleles coalesce (entry 3 of row 1) if they are either descended from the same allele, or if gene conversion acted in the parent. Otherwise, if they are descended from different alleles from the same parent, then they remain distinct if gene conversion does not act (entry 2 of row 1). The frequency of sex *σ* does not affect the terms in row 1, as the probabilities of identity by descent from a single parent are the same under both sexual and asexual reproduction, if we assume that unpaired alleles are equally likely to be sampled from one of the two allele copies. These probabilities could change if there was biased sampling of alleles when sex is rare; I will discuss this point when analysing the ‘very rare sex’ regime.

**Figure 1.**
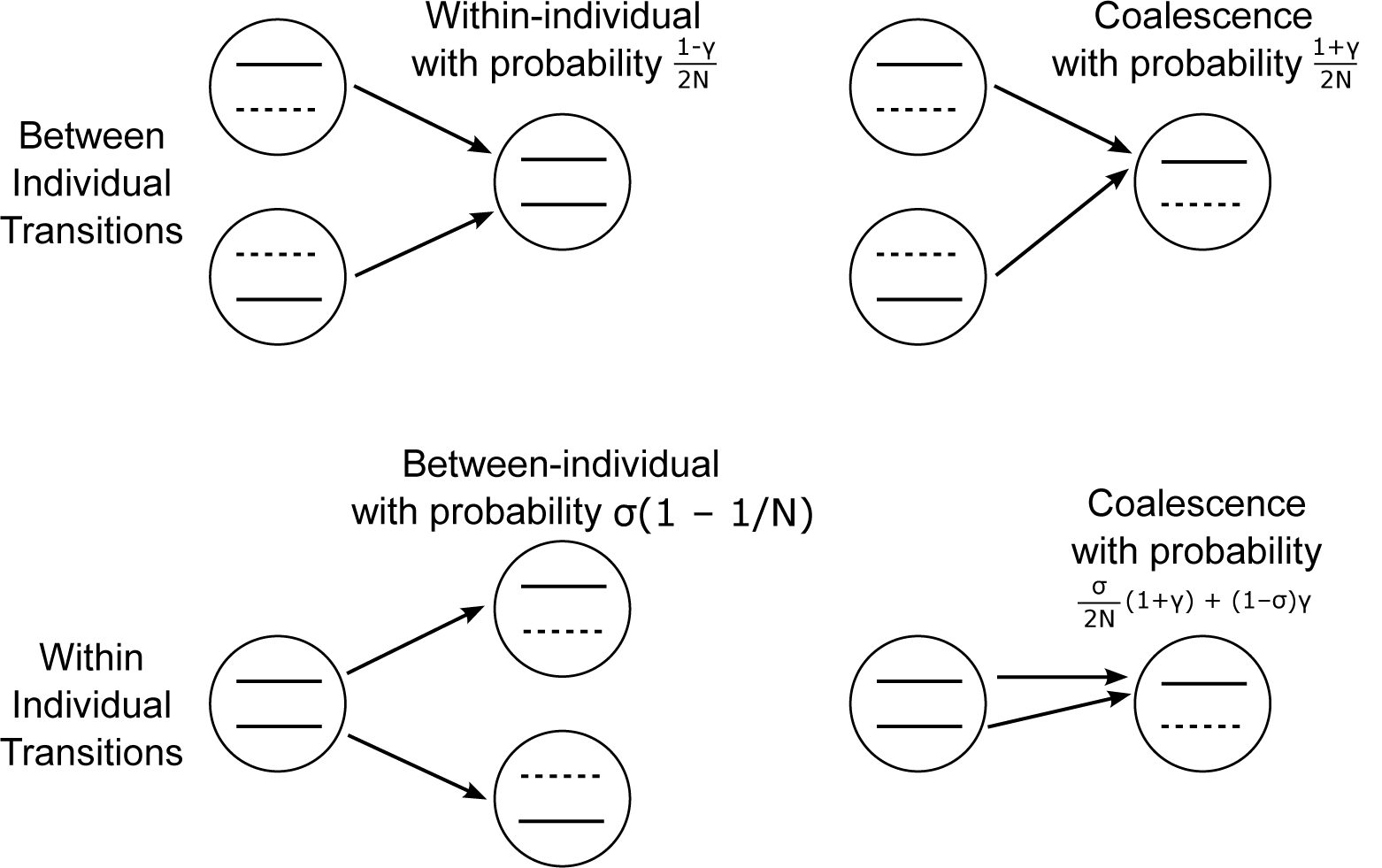
Schematic of the transition probabilities in the facultative sex coalescent. Sampled alleles are shown as solid lines, while dashed lines are alternate alleles that are not sampled. Figure is originally from Hartfield *et al*. (2016) and is reprinted with permission from the Genetics Society of America.

If the two alleles are taken from different genetic backgrounds within an individual, they coalesce (entry 3 of row 2) if there is either sexual reproduction followed by inheritance from the same parent, or gene conversion in the absence of sex. They can also be descended from distinct alleles in separate individuals if the offspring was created by sex involving two distinct parents (entry 1 of row 2). The diagonal entries are one minus the other entries in each row. Hartfield *et al*. (2016) contains further details on how the transition probabilities are formed.

### Simulations

Analytical results for pairwise coalescent times will be compared to stochastic simulations written in C, which are based on those used in Agrawal and Hartfield (2016). The simulation tracks a single neutral bi–allelic locus in a facultative sexual population forwards in time. Each generation, a proportion *σ* of reproductions are sexual, with offspring genotypes generated according to Hardy–Weinburg equilibrium frequencies. The remaining fraction 1 − *σ* of reproductions are as asexual clones. Mitotic gene conversion acts with probability *γ*, which converts heterozygotes to homozygotes with equal probability (i.e., gene conversion is unbiased). Using these deterministic expectations, the number of genotypes among *N* individuals is drawn from a multinomial distribution to implement random drift. Neutral mutations are sequentially introduced, each time from a single copy. The pairwise diversity *x*(1 − *x*) (for *x* the derived allele frequency) is summed over the neutral allele trajectory, until the mutation is either fixed or lost. Ten million neutral alleles are introduced and their summed pairwise diversity values calculated; the mean over all introductions equals the coalescent time, scaled to that expected for the standard coalescent (Charlesworth *et al*. 1993; Nordborg *et al*. 1996). Confidence intervals are calculated from 1,000 bootstraps.

## Results

### Approximate coalescent times for two alleles

I will first look at two–allele results to determine the long–term pairwise coalescent time, then subsequently determine if a coalescent *N*_*e*_ can be defined in each case. I will also relate two–allele results to *F* −statistics (Wright 1951) under each scenario. Results can be summarised by three phases, which depend on the relative frequencies of sex and gene conversion compared to the actual population size, as shown in Figure 2):

**Figure 2.**
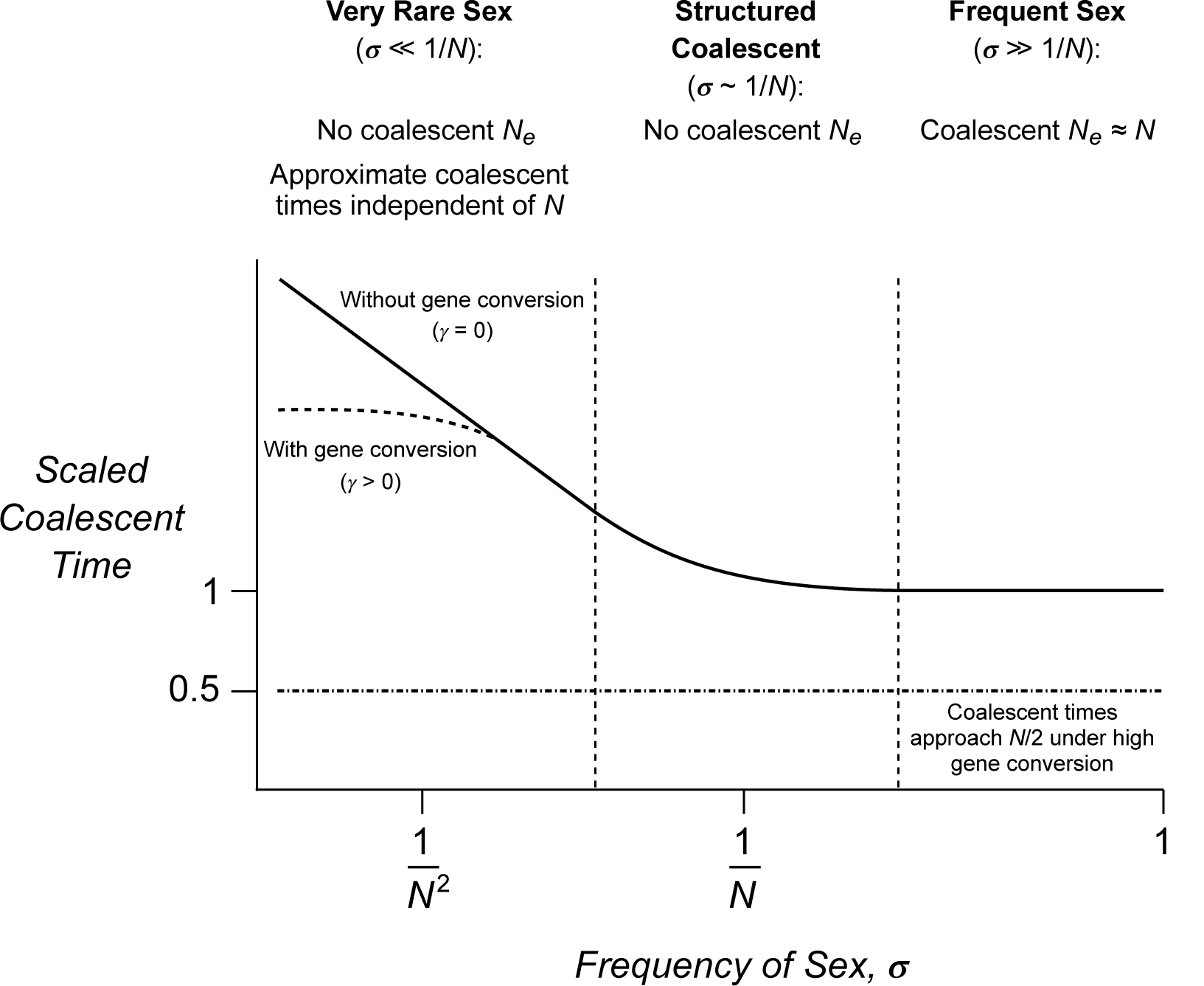
Outline of scaled coalescent times for two alleles under facultative sex. Note that the results given at the top of the figure assume that gene conversion *γ* acts on the same order as sexual reproduction. If gene conversion is much more frequent, then coalescent times tend to *N/*2 as shown by the dotted–dashed line.

1. **The ‘frequent sex’ regime** (*σ* ≫ 1*/N*).
  a. *If gene conversion is rare* (*γ* ≪ 1), then due to the high occurrence of genetic segregation the resulting coalescent process is similar to the standard coalescent, and the coalescent *N*_*e*_ ≈ *N*.
  b. *If gene conversion is high* (*γ* → 1), coalescent *N*_*e*_ = *N/*2 as heterozygosity is removed.
2. **The ‘structured coalescent’ regime** (*σ* ∼ 1*/N*).
  a. *If gene conversion also acts with probability* ∼1*/N*, state transitions (i.e., whether the two alleles lie in the same or different individuals) and coalescent events occur at the same relative frequencies. Hence, population history cannot be captured by a coalescent *N*_*e*_.
  b. *If gene conversion is high* (*γ* ≫ 1*/N*), coalescent *N*_*e*_ = *N/*2, similar to the ‘frequent sex’ regime.
3. **The ‘very rare sex’ regime** (*σ* ≪ 1*/N*).
  a. *If gene conversion is also very weak* (*γ* ≪ 1*/N*) then Möhle’s theorem can be used to derive approximate two–allele coalescent times, which only depend on the frequency of sex and gene conversion and are independent of *N*. These times do not translate into a coalescent *N*_*e*_.
  b. *If gene conversion is much more frequent than sex* (*γ* ≫ 1*/N*), then either no coalescent *N*_*e*_ exists (if *γ* ∼ 1*/N*) or *N*_*e*_ = *N/*2 (if *γ* ≫ 1*/N*). Simulations suggest that the scaled coalescent time is halved in most cases.

The ‘structured coalescent’ regime was previously outlined in Hartfield *et al*. (2016) so will not be discussed here. I will instead elucidate the coalescent *N*_*e*_ when sex is frequent, and introduce results for the ‘very rare sex’ regime.

### The ‘frequent sex’ regime

As a first example of how Möhle’s theorem provides insight into how the facultative sex coalescent can be approximated, I partition 𝕋 with respect to the scaling factor *ϵ* = 1/2*N* to determine the fast and slow–rate events. Since I do not make any further assumptions on the frequencies of sex and gene conversion, then the ensuing result applies when both these events are frequent (more precisely, when *σ, γ* ≫ 1*/N*). The two–allele results were previously presented in the supplementary matierial of Hartfield *et al*. (2016); here I show how they can be used to define a coalescent *N*_*e*_ for a genealogy of any size. More details on all matrix calculations are available in Supplementary File S1.

𝕋 can be written as 𝔸 + 𝔹/2*N*, with the sub–matrices defined as:

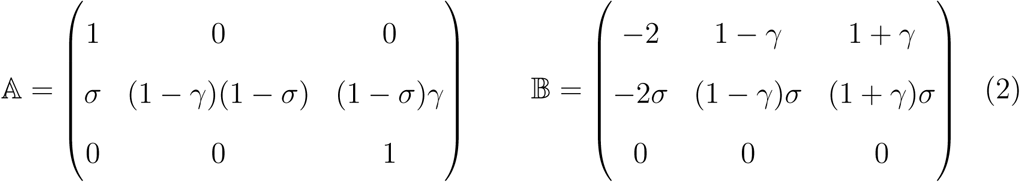

Using Möhle’s theorem, the short–term matrix ℙ and long–term matrix 𝔾 equal:

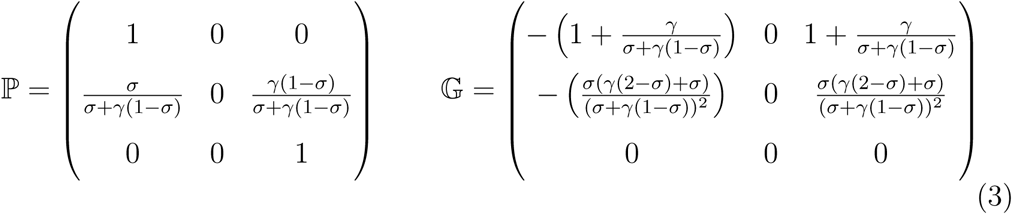

A potential coalescent effective population size *N*_*e*_ is inferred by inspecting ℙ and 𝔾. ℙ shows that over short timescales (much less than 2*N* generations in the past), alleles will either segregate into different individuals with probability 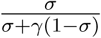 or coalesce with probability 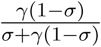. 𝔾 implies that, if alleles have not coalesced, they do so over the long term with an increased rate of 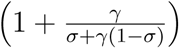 per 2*N* generations. For a Wright–Fisher population, the coalescent timescale is obtained in the standard model by scaling time by 2*N*, so any two alleles coalesce at rate 1 per coalescent generation. Under this approximation, the standard coalescence rate is obtained by rescaling time by 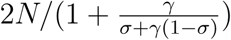.

To determine if this rescaling does indeed constitute a coalescent *N*_*e*_, it needs to be shown that it causes a genealogy of any size to converge to the standard coalescent. The short–term matrix ℙ (Equation 3) shows that each pair of alleles from the same individual will quickly segregate out into different individuals, or coalesce (the latter being unlikely in the biologically realistic case of *γ* ≪ *σ*). Let there be *n* alleles remaining in different individuals after this readjustment. The transition matrix of the subsequent coalescent process is modelled using three states: (1) *n* alleles are present in *n* distinct individuals; (2) *n* alleles are present in *n* − 1 distinct individuals; (3) there is a coalescent event. Because sex is so frequent, I further assume that it is unlikely that *n* alleles will be present in less than *n* − 1 individuals in a single generation. The model only considers the genealogical history up to the first coalescent event. The transition matrix is the same as for the two–allele case (Equation 1) except that the first row now equals:

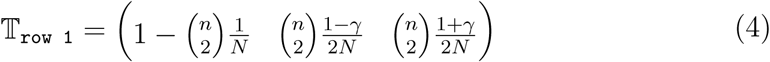

Applying Möhle’s theorem with the long–term matrix scaled by 1/2*N* gives the same ℙ (Equation 3), but 𝔾 is multiplied by a factor 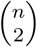. Hence after rescaling time by 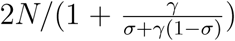, the coalescent rate equals 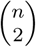 as with the standard coalescent. Hence, a coalescent *N*_*e*_ can be defined and is equal to 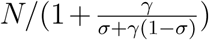.

This result is a similar form to the general reduction in *N*_*e*_ = *N/*(1 + *F*) obtained under various forms of inbreeding (Caballero and Hill 1992), with *F* equal to 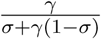. If the probability of gene conversion is low relative to the frequency of sex (i.e., *γ* ≪ 1) then *F* ≈ *γ/σ* ≪ 1, so *N*_*e*_ ≈ *N*. *F* increases with *γ* up to its maximum value of 1 when *γ* = 1. It therefore follows that the coalescent *N*_*e*_ = *N/*2 under this scenario, due to immediate within–individual coalescence. In practice, such a drastic reduction in *N*_*e*_ is unlikely given the low probability of gene conversion affecting a single site. For example, Sharp and Agrawal (2016) estimated a mitotic gene conversion frequency of ∼10^−6^ per basepair per generation in *Drosophila melanogaster*.

### The ‘very rare sex’ regime

Möhle’s theorem can also be applied when the frequency of sex is extremely low relative to the population size (*σ* ≪ 1*/N*). Here, the slow–rate matrix is scaled by a parameter different from the population size. I will assume both rare sex and gene conversion (i.e., *σ, γ* ≪ 1*/N*) and use *λ* = *σ* +*γ* to determine the slow–rate matrix. It is also convenient to make the substitution *ϕ* = *σ/γ*, which determines whether diploid genotypes experience allelic sequence divergence (*ϕ* > 1) or convergence due to gene conversion (*ϕ* < 1) (Hartfield *et al*. 2016).

After transforming *σ, γ* into their new variables and removing terms of 𝒪(*λ*^2^), the transition matrix 𝕋 can be written as 𝔸 + *λ*𝔹 + *o*(*λ*^2^), with the sub–matrices equal to:

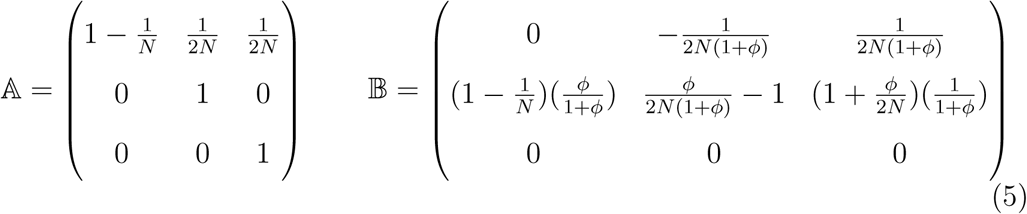

Applying Möhle’s theorem to obtain ℙ, 𝔾:

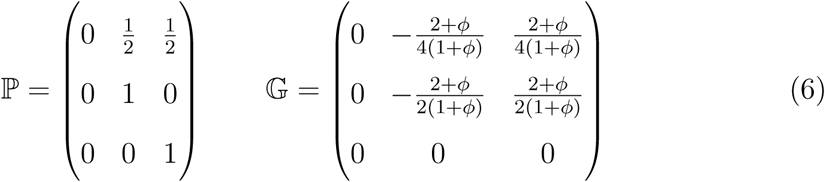

Here, the short–term matrix ℙ shows that two alleles in different individuals will either segregate into the same individual to become a set of paired alleles, or coalesce. Either event is equally likely to occur. Two alleles from the same individual will remain as such, so it is the only remaining state. The long–term matrix 𝔾 further shows that a set of paired alleles will coalesce at rate 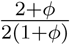 per 1*/λ* generations. Hence, if discrete time is scaled by 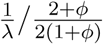 then a within– individual allele pair will coalesce at rate 1 per rescaled time unit. Restoring back the *σ, γ* terms gives the expected within–individual coalescent time:

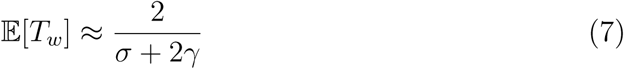

The key result here is that under very rare frequencies of sex, the two–allele approximate coalescent time is independent of the actual population size *N*. The exact coalescent time is affected by *N* (Hartfield *et al*. 2016), but in this regime coalescent times are most strongly influenced by the rare occurrences of sex and gene conversion, which reduces the probability that two alleles will meet their common ancestor. It is possible to re–write Equation 7 using the compound parameters Ω = 2*Nσ*, Γ = 2*Nγ* to derive 𝔼 [*τ*_*w*_], the mean coalescent time on the coalescent timescale (that is, time is scaled by 2*N*):

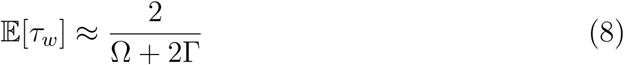

𝔼 [*τ*_*b*_] is simply half of 𝔼 [*τ*_*w*_]. This is because if two alleles are sampled from different individuals, then the long–term state will only be entered with probability 1/2 (see ℙ in Equation 6), otherwise the two alleles will coalesce ‘instantaneously’ (more specifically, on a timescale much less than 𝒪(*λ*)). Note that 𝔼 [*τ*_*w*_], 𝔼 [*τ*_*b*_] as given here are equivalent to Equation 11 in Hartfield *et al*. (2016) but if only retaining the second fraction term that is of 𝒪(*λ*), as rare sex and gene conversion most strongly influence the expected coalescent time.

Here too, the scaled coalescent time can be related to the inbreeding coefficient *F*. Recall that *F* ranges between −1 and 1; negative values denote an excess of heterozygosity, while positive values indicate a heterozygote deficit (Wright 1951). By comparing the within–individual coalescent time 𝔼 [*τ*_*w*_] to the general term 1/(1 + *F*), the two equate if *F* = (2Γ + Ω − 2)/2 (Figure 3). If Ω < 2(1 − Γ) then *F* is negative, as sex is sufficiently rare to cause some degree of allelic sequence divergence, increasing heterozygosity. *F* reaches its minimum of −1 when both Ω, Γ are zero. Otherwise, *F* is positive as gene conversion removes heterozygous sites, with a maximum of *F* = 1 attained when Γ = (4 − Ω)/2. As *F* cannot exceed one, then this bound implies an upper limit to Ω, Γ at which the rare–sex approximations are valid.

**Figure 3.**
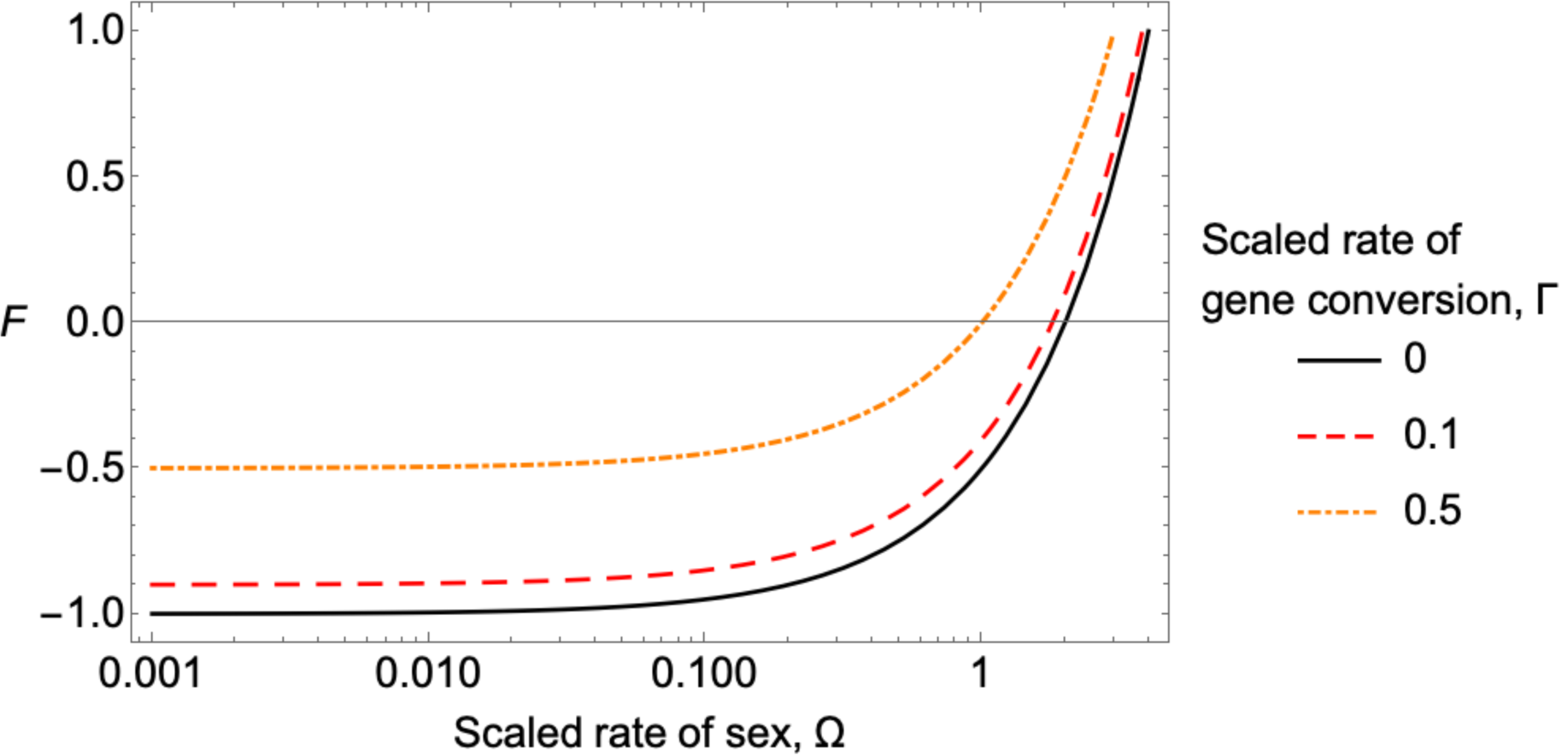
Inbreeding coefficient *F* under rare sex, based on the within–individual coalescent time. Plot of *F* = (2Γ + Ω − 2)/2 as a function of Ω = 2*Nσ*. Different lines represent different Γ = 2*Nγ* values as shown in the legend.

Although an approximate coalescent time for two alleles can be obtained, can it be defined as a coalescent *N*_*e*_? The answer is no, as the pairwise coalescence rate is altered as more alleles are introduced. To provide a counterexample, I outline a transition matrix for the case of two sets of alleles from two individuals, so there are four alleles in total. There are five states representing all different partitions of these alleles among individuals (i.e., two paired alleles; one paired and two unpaired alleles; four unpaired alleles), and possible coalescent events (either one or two alleles coalesce in a generation). This model only determines the process until the first coalescence event.

The transition matrix under this scenario is outlined in Appendix A. As before, Möhle’s theorem is applied with *λ* = *σ* + *γ* determining long–term events. The short–term matrix ℙ becomes:

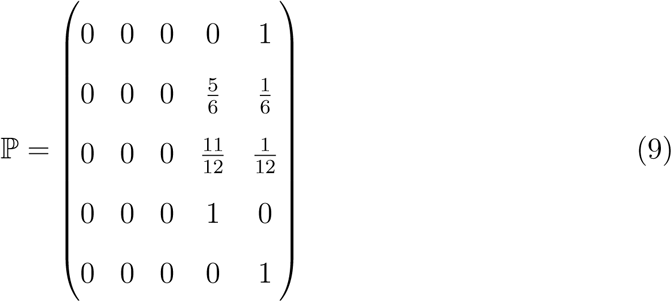

and 𝔾 = 0. Hence, there will be at least one coalescent event over short timescales (states 4 and 5 represent single and double coalescent events, respectively). In particular, if there are initially two sets of paired alleles then they will coalesce into a single pair by the end of the initial phase (as given in row 1). Unlike the standard coalescent, these coalescent events do not occur at regular times proportional to 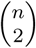, but much more frequently. Heuristically, this property arises since individuals (and hence pairs of alleles) coalesce with probability 𝒪(1*/N*), but the final two alleles coalesce with probability 𝒪(*λ*) ∼ 𝒪(1*/N* ^2^). This process creates a genealogy with very short terminal branches and two long internal branches representing divergence of the two remaining alleles (see Figure 5 of Hartfield *et al*. (2016)). Thus, unlike the frequent–sex case, a coalescent *N*_*e*_ cannot be defined.

Further analysis of the *n* > 2 case is desirable to determine how the coalescent process transitions from the ‘fast’ state, where events occur over 𝒪(*N*) generations, to the ‘slow’ state when two alleles remain and coalesce over 𝒪(*N* ^2^) generations. To do so, I approximate the transition matrix to only focus on 𝒪(1*/N*) events in the fast state (i.e., coalescent events that do not involve sex or gene conversion). Let there be *n* = 2*x* + *y* alleles, of which *x* are paired alleles that are sampled from the same individual, and *y* are unpaired alleles. The maximum number of paired alleles *x*_*m*_ equals the largest whole number that is less than or equal to *n/*2 (i.e., [*n/*2] in mathematical notation). It is possible to define a square transition matrix 𝕋_*n*_ with *x*_*m*_ +3 rows and columns. The first *x*_*m*_ +1 rows denote states where there are *x*_*m*_, *x*_*m*_ − 1 … 1, 0 set of paired alleles; row *x*_*m*_ + 2 the absorbing state caused by a single coalescent event, and row *x*_*m*_ + 3 the absorbing state caused by a double coalescent event. The entries of 𝕋_*n*_ are given in Appendix B.

The transition to the slow state depends on the order of single to double coalescent events before two alleles remain. 𝕋_*n*_ can be written in the canonical form for Markov chains (Grinstead and Snell 1997) to determine how much time is spent in the fast state, depending on how alleles were initially sampled:

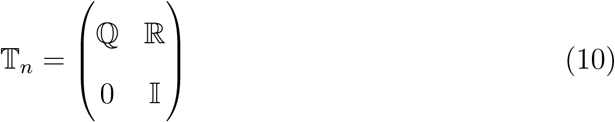

ℚ is a (*x*_*m*_ + 1) × (*x*_*m*_ + 1) matrix of non-coalescent states, ℝ is a (*x*_*m*_ + 1) × 2 matrix denoting transition to coalescent states, and 𝕀 a 2×2 identity matrix. From this form, we can subsequently derive ℕ = (𝕀 − ℚ)^−1^ which denotes the expected time spent in each non–coalescent state before a coalescent event occurs. ℕ ℝ is the probability of ending up in each coalescent state.

Appendix B provides example calculations when there are three or four alleles in the tree. In summary, the fast–state is shortest with three alleles if they are all sampled from different individuals, while with four alleles the fast–state is shortest when two sets of paired alleles are sampled. This latter results arises because coalescence is more likely with just paired alleles (occurring with probability 𝒪(1*/N*), rather than 𝒪(1/2*N*) with unpaired alleles). Sampling just paired alleles will more efficiently capture the effects of polymorphism as shaped by rare sex and gene conversion, and minimise the confounding influence of recent mutation.

Note that these results are based on the assumption that unpaired alleles are randomly sampled from one of the two possible alleles in diploids. The results would differ if biased sampling were to occur. If phased genome data were available, then it would be possible to instead sample one of the two diverged alleles per individual. If sex and gene conversion were negligible in the recent past, then these unpaired alleles will follow a standard coalescent process with coalescent probability proportional to 1*/N* [as opposed to 1/2*N* under the previous assumptions; see also Ceplitis (2003)], hence the coalescent *N*_*e*_ = *N/*2. This sampling procedure can inform on mutations appearing 𝒪(*N*) generations ago, but not on how ancient sex and gene conversion events shape within–individual polymorphism.

### Simulation Comparisons of Möhle’s approximations

Figure 4 plots the two–allele scaled coalescent time (specifically, the between– individual time 𝔼 [*τ*_*b*_] for the frequent sex and very rare sex cases) as compared to simulations. Results are provided for different values of *σ*, the frequency of sexual reproduction, and Γ = 2*Nγ*, the population–scaled gene conversion rate. Analytical results are generally accurate for low gene conversion rates (Γ ≤ 0.5), but become underestimates as Γ approaches 0.5 (Figure 4a). Analytical solutions also underestimate the scaled coalescent time if *σ* ∼ 10^−5^ (equivalent to Ω = 2*Nσ* ∼ 1 for the population size used in simulations), as the ‘structured coalescent’ regime is entered. These results exemplify how rare sex can substantially inflate coalescent times. For example, if Ω = 0.001 then the scaled coalescent time is 1,000–fold larger than in the standard coalescent, in the absence of gene conversion. As Ω exceeds 1 then coalescent time approximate to those in the standard coalescent.

**Figure 4.**
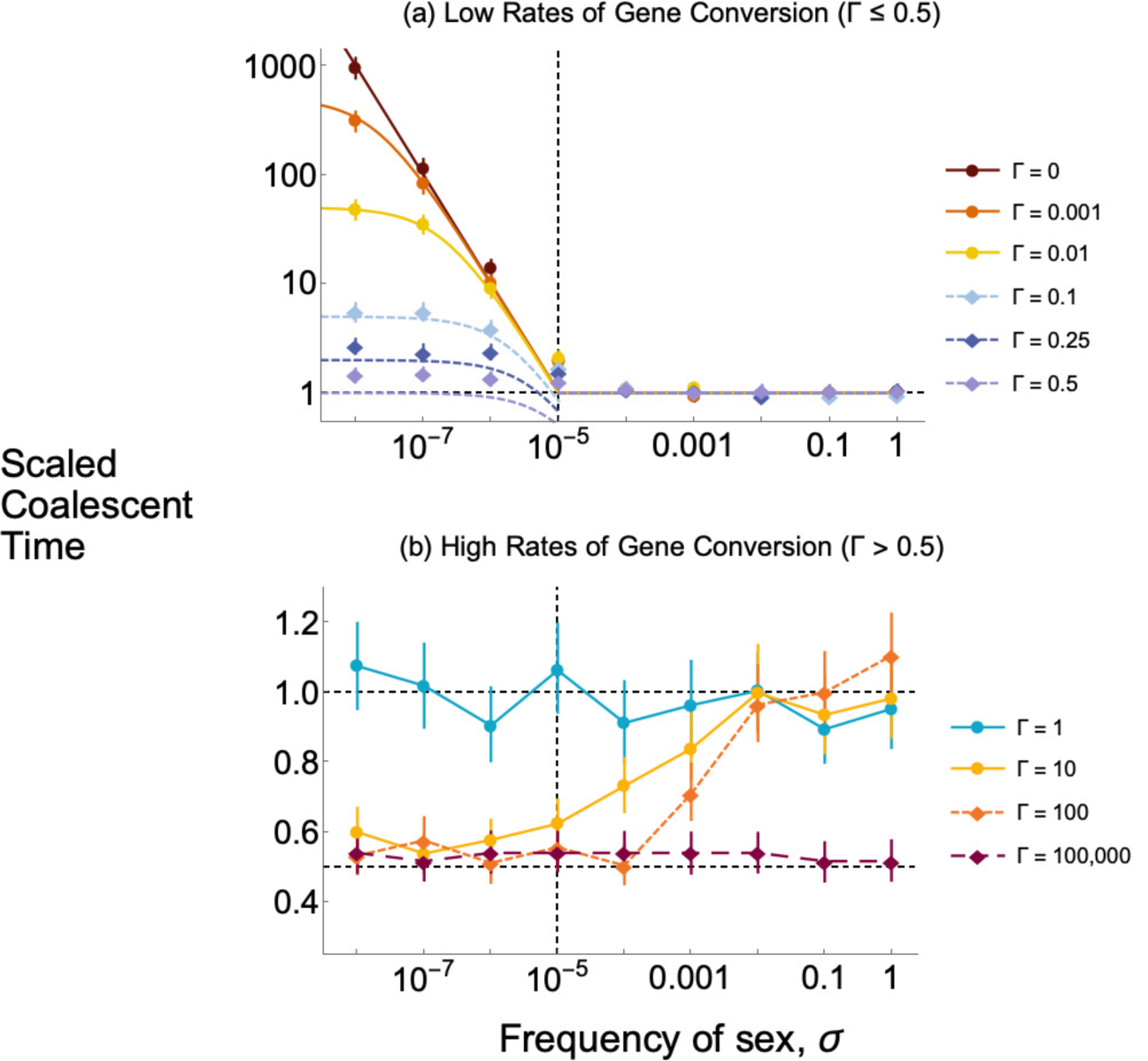
Simulation comparisons of scaled between–individual mean coalescent time. Plots of the scaled coalescent time as a function of the frequency of sex *σ*. Colours represent different rates of gene conversion, as shown in each subplot legend. Simulations assume a diploid population of size *N* = 50,000. Points are simulation results, with bars representing 95% confidence intervals for the mean value (if they cannot be seen, they completely lie within the point). (a) Results for low rates of gene conversion (Γ ≤ 0.5). Solid lines are analytical approximations (1 for Ω > 1, corresponding to *σ* > 10^−5^, as shown by the vertical dashed line; 1/(Ω + 2Γ) for Ω < 1). Horizontal dashed line denote a scaled time of 1. (b) Results for high frequencies of gene conversion (Γ > 0.5). Horizontal dashed lines show scaled times of 0.5 and 1; vertical dashed line denotes *σ* = 10^−5^. Note the different *y* − axis scales in each subplot. Simulation results are also available in Supplementary File S1.

For very high frequencies of gene conversion (Γ > 0.5; Figure 4b) coalescent times are generally on the same order as the standard coalescent. As Γ increases then the coalescent times are half that in the standard coalescent, due to gene conversion causing rapid within–individual coalescence (biologically, this mechanism manifests itself through reduced heterozygosity). The amount of gene conversion needed for this halving to occur depends on the frequency of sex. For example, coalescent times are nearly halved if Γ = 10 for Ω = 0.001, but an extremely large value of Γ = 100, 000 is required in obligate sexual populations.

## Discussion

Here, I have outlined calculations to determine how to approximate the coalescent under facultative sex, with an emphasis on determining when it converges to the standard coalescent. I first determined approximate pairwise coalescent times using Möhle theorem, for cases where sex is frequent or very rare (Figures 2, 4). I then further determined if a coalescent *N*_*e*_ can be subsequently defined. If sex is frequent (*σ* ≫ 1*/N*), then a coalescent *N*_*e*_ exists and approximates *N*. For very rare sex (*σ* ≪ 1*/N*), pairwise coalescent times can be inflated due to allelic sequence divergence, the extent of which is effectively independent of the actual population size. However, a coalescent *N*_*e*_ does not exist due to coalescence acting on a much faster timescale when there are more than two alleles. I subsequently analysed the coalescent process with more than two alleles, to determine how the initial allele sample would affect the transitions from this ‘fast’ state, to the ‘slow’ state when only two alleles remain in the tree.

The reasons why a coalescent *N*_*e*_ does not exist under very rare sex is exemplified by how the use of Möhle’s theorem differs from that in most previous studies. In other cases where Möhle’s theorem is used to approximate coalescent models (e.g., population structure or self–fertilisation), there is first rapid coalescence within a group [e.g., within a subpopulation under an island model with weak migration (Wakeley 2004), or within individuals under self–fertilisation (Nordborg and Donnelly 1997; Nordborg and Krone 2002)]. Coalescence then occurs under a rescaled standard coalescent among remaining alleles between groups. Under very rare sex with weak gene conversion, the opposite behaviour arises; there is first a coalescence of alleles from different individuals, followed by extended coalescent times for a pair of alleles within individuals.

How can genome data from facultative sexuals be best analysed? If sex is frequent then the coalescent process is similar to the standard coalescent, with a slightly adjusted *N*_*e*_. It has also been previously shown that the degree to which linkage disequilibrium is broken down by meiotic recombination also scales with the frequency of sex, if it is not rare (Hartfield *et al*. 2018). Together, these results suggest that for species with facultative but frequent sex, using models based on the standard coalescent would well approximate their gene genealogies following appropriate rescaling of recombination rates. However, once sex becomes rare and individuals start exhibiting allelic divergence [as proposed for, e.g., the human pathogen *Trypanosoma brucei gambiense* (Weir *et al*. 2016)], then a coalescent *N*_*e*_ cannot be defined. In this scenario, analysing different combinations of alleles will inform on either the historical or more recent forces shaping diversity. When four alleles are sampled (Appendix B), the fast–state is minimised in a tree with two sets of paired alleles, which will provide most information on how rare sex and gene conversion have shaped within–individual allele divergence. Conversely, analysing a tree composed of the same type of diverge allele but taken from different individuals will provide information on recent mutations (‘recent’ meaning arising approximately *N* generations ago). Methods using the distribution of coalescent times from a small number of alleles (e.g., the generating–function method of Lohse *et al*. (2011)) could be particularly useful to determine how historical frequencies of sex and gene conversion have shaped genetic diversity. However, these results chiefly apply when sex is sufficiently infrequent so that allele divergence could occur; these results break down if gene conversion was common, or if the facultative sexual species was recently–derived from an obligate sexual ancestor. For more complex scenarios, it may be necessary to use bespoke coalescent models and simulation software that explicitly consider facultative sex (Hartfield *et al*. 2016, 2018).

The results presented here only consider the effect of neutral processes on *N*_*e*_ and coalescent histories, and can change in the presence of selection. For example, background selection reduces local *N*_*e*_ (and hence local diversity) (Charlesworth *et al*. 1993; Hudson 1994); its effects can be amplified in facultative sexuals due to both a lack of both recombination and segregation (Agrawal and Hartfield 2016). These calculations also do not include population or life–stage structure (Orive 1993). However, the effects of population structure can be easily incorporated into the transition matrix (Hartfield *et al*. 2016), and coalescent approximations can also be determined under different cases of the island model (Wakeley 2004).

Overall, these calculations clarify how facultative sex affects the genealogical history of a population, and when it can be approximated by the standard coalescent. They can also be used in future modelling work of evolution under facultative sex, to determine both how neutral genetic diversity is affected, and how to best analyse and interpret genome data in other scenarios.

## Acknowledgments

I would like to thank Maria Orive for inviting me to speak at the 2019 AGA Presidential Symposium “Sex and Asex: The genetics of complex life histories”, and Derek Setter, Konrad Lohse, Brian Charlesworth, Maria Orive and two anonymous reviewers for providing comments on the manuscript.

## Data availability

Simulation code, and Supplementary File S1, is available from http://github.com/MattHartfield/FacSexNe.

## Appendix A Very rare sex with four alleles

Here, I outline the exact coalescent process for four alleles. The underlying transition matrix has five states: (1) two sets of paired alleles; (2) one set of paired alleles, and two unpaired alleles; (3) four unpaired alleles; (4) the number of alleles is reduced by one due to a single coalescent event; (5) the number of alleles is reduced by two due to two paired alleles coalescing (i.e., both pairs are descended from the same individual).

The transition probabilities for each state are as follows. Note that each probability is considered up to 𝒪(1*/N* ^3^).

From state 1:

- To state 2: occurs due to sex with probability 2*σ* (note the factor of two due to two sets of paired alleles).
- To state 3: requires multiple sex events of order *σ*^2^ = 𝒪(1*/N* ^4^).
- To state 4: occurs due to gene conversion within one of the paired alleles, with probability 2*γ*.
- To state 5: requires the two individuals to coalesce with probability 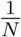.

From state 2:

- To state 1: Requires that (i) the two unpaired alleles to descend from different alleles within the same individual, and (ii) the paired allele descends from a different individual. The probability is 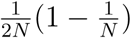.
- To state 3: Requires the paired allele to segregate by sex, with probability *σ*.
- To state 4: Requires either (i) the two unpaired alleles to coalesce, and the paired allele to descend from a different individual, with probability 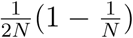; (ii) 1 unpaired allele coalescing with the paired individual with probability 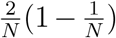; (iii) the paired allele experiences gene conversion with probability *γ*. The total probability is 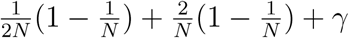.
- To state 5: Requires all three individuals to be descended from the same parent with probability 1*/N* ^2^.

From state 3:

- To state 1: Requires all four alleles to place themselves in two different individuals, with each allele having a unique descendant. There are 3 possible set of two pairs. The overall probability is 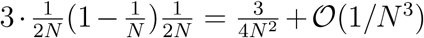.
- To state 2: Requires two alleles to descend from the same parent, and other two from different parents. The total probability is 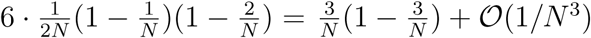.
- To state 4: Requires two alleles to coalesce, and the other two to descend from different alleles with probability 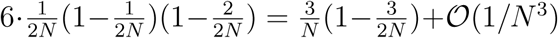.
- To state 5: Requires two individual coalescent events with probability 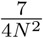 [there are seven ways in which two alleles can coalesce from four alleles; see Wakeley (2009, Equation 3.12)].

States 4 and 5 absorbing.

Hence the transition matrix 𝕋 = 𝔸 + *λ*𝔹 with:

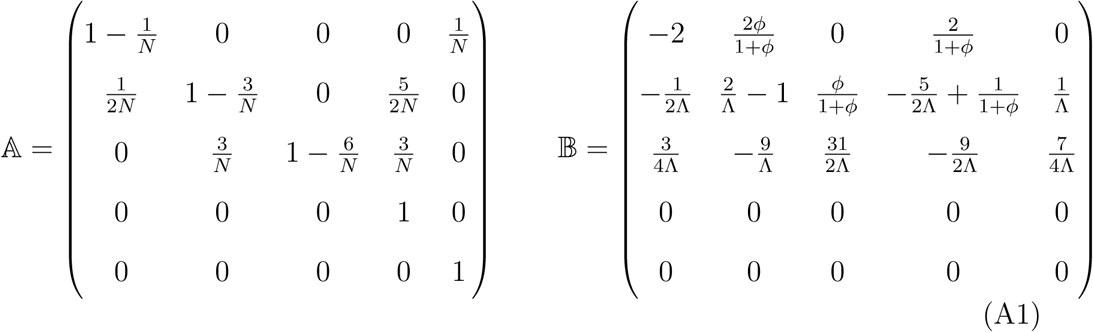

where Λ = *N* ^2^*λ*. Applying Möhle’s theorem gives ℙ as given in Equation 9.

## Appendix B Approximation of the very rare sex regime

### General approach for *n* alleles

The fast–state transition matrix 𝕋_*n*_ contains the following 𝒪(1*/N*) transition probabilities (see also Table 1 of Hartfield *et al*. (2016)), with 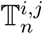 denoting the entry in row *i* and column *j*:

- 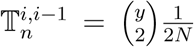 is the probability that two unpaired alleles form a paired sample.
- 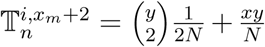 is the probability that a single coalescent event occurs.
- 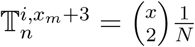 is the probability that two paired alleles will coalesce.
- 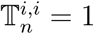 minus the sum of the probabilities listed above.

#### Example with three and four alleles

We can demonstrate the utility of the above method by analysing results with three and four alleles, and combining results to determine how the resulting coalescent tree differs depending on how alleles were sampled.

With three alleles, there are only three states: (i) one paired sample and one unpaired sample, (ii) three unpaired samples, (iii) single coalescence event. Note there is only one coalescent state; a double coalescence is not possible as there can only be one set of paired alleles. The transitions matrix is:

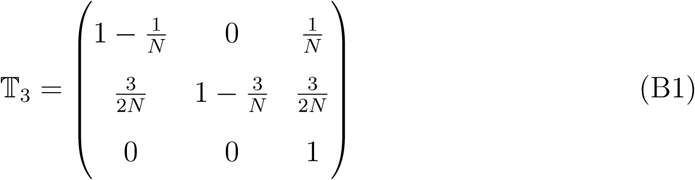

Using the canonical form of 𝕋_*n*_ in Equation 10, we can find the matrix N of mean time spent in each state before coalescence (with each result scaled by 2*N* to be on the coalescent timescale):

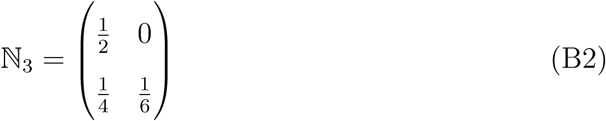

If ℕ is multiplied by (1, 1)^*T*^ where *T* denotes a transpose, then we obtain the mean time until coalescence for each initial state (Slatkin 1991), which equals (1/2, 5/12). Hence, the time to reach the slow state is shorter when three unpaired alleles are taken.

For four alleles, I approximate the exact transition matrix by focussing on 𝒪(1*/N*) events, which gives 𝔸 in Equation A1. By putting this matrix into the canonical form of Equation 10, it is possible to obtain the mean coalescent times for each starting configuration, which are (1/2, 1/4, 5/24). The product ℕℝ denoting the probability of ending up in each coalescent state is:

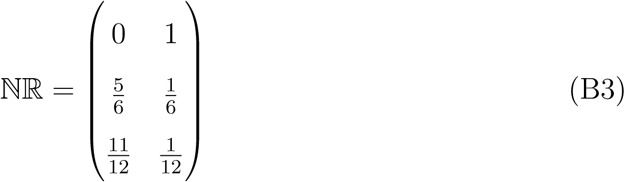

Note that this matrix is equal to the 3 × 2 top–right corner of the fast–matrix ℙ (Equation 9).

We can combine results from the three–allele and four–allele matrix to determine the structure of coalescent trees depending on how alleles are sampled.

- With two paired alleles, they will undergo a double coalescent event after an average of 1/2 coalescent generations (equivalent to *N* discrete generations), at which point the process will enter the slow state.
- With one set of paired alleles and two unpaired alleles, a coalescent event arises after an average of 1/4 generations. Equation B3 states that with probability 1/6, this will be a double coalescent event, whereas with probability 5/6 there will only be a single coalescent event. If so then the three–allele process will then start, with a configuration of one set of paired alleles and one unpaired allele. There will then be an additional 1/2 generations on average before another coalescent event, starting the slow state. Hence, there will be an average of 1/4 + 5/6 · 1/2 = 2/3 coalescent generations in the fast state.
- With four unpaired alleles, a coalescent event occurs after an average of 5/24 generations. A double coalescent event occurs with probability 1/12 instantly triggering the slow state, otherwise the three–allele process starts with one set of paired alleles and one unpaired allele. It then takes an average of 5/12 generations to enter the slow phase. Hence the mean time of the fast phase is 5/24 + 11/12 · 5/12 = 85/144 ≈ 0.59 generations.

